# Ultra-High Dose Rate Helium Ion Beams: Minimizing Brain Tissue Damage while Preserving Tumor Control

**DOI:** 10.1101/2024.06.13.598785

**Authors:** Ivana Dokic, Mahmoud Moustafa, Thomas Tessonnier, Sarah Meister, Federica Ciamarone, Mahdi Akbarpour, Damir Krunic, Thomas Haberer, Jürgen Debus, Andrea Mairani, Amir Abdollahi

## Abstract

Ultra-high dose rate radiotherapy with electrons and protons has shown potential for cancer treatment by effectively targeting tumors while sparing healthy tissues (FLASH effect). This study aimed to investigate the potential FLASH sparing effect of ultra-high-dose rate helium ion irradiation, focusing on acute brain injury and subcutaneous tumor response in a preclinical in vivo setting. Raster-scanned helium ion beams were used to compare the effects of standard dose rate (SDR at 0.2 Gy/s) and FLASH (at 141 Gy/s) radiotherapy on healthy brain tissue. Irradiation-induced brain injury was studied in C57BL/6 mice via DNA damage response, using nuclear γH2AX as a marker for double-strand breaks (DSB). The integrity of neurovascular and immune compartments was assessed through CD31^+^ microvascular density and activation of microglia/macrophages. Iba1+ ramified and CD68^+^ phagocytic microglia/macrophages were quantified, along with the expression of inducible nitric oxide synthetase (iNOS). Tumor response to SDR (0.2 Gy/s) and FLASH (250 Gy/s) radiotherapy was evaluated in A549 carcinoma model, using tumor volume and Kaplan-Meier survival as endpoints. The results showed that helium FLASH radiotherapy significantly reduced acute brain tissue injury compared to SDR, evidenced by lower levels of DSB and preserved neurovascular endothelium. Additionally, FLASH radiotherapy reduced neuroinflammatory signals compared to SDR, as indicated by fewer CD68+ iNOS+ microglia/macrophages. FLASH radiotherapy achieved tumor control comparable to that of SDR radiotherapy. This study is the first to report the FLASH sparing effect of raster scanning helium ion radiotherapy in vivo, highlighting its potential for neuroprotection and effective tumor control.

## Introduction

Helium ion radiotherapy provides exceptional precision in targeting tumors due to the Bragg peak phenomenon and reduced lateral scattering. This modality has gained prominence in the radiation oncology community because of its potential to minimize damage to healthy surrounding tissues despite conformal coverage of the tumor volume [1]. Moreover, helium ions are characterized by a higher linear energy transfer (LET) compared to e.g. protons. Irradiation with higher LET induces complex DNA damage that is more challenging to repair, leading to an increased RBE and potentially enhancing both tumor control and therapeutic efficacy. Helium ions are particularly noteworthy for their significantly reduced lateral scattering, which is approximately 50% smaller than that observed with protons [2]. Helium ions are being actively explored as a potential alternative to proton therapy, with in silico indications suggesting potentially better outcomes under certain clinical scenarios [3]. The risk of healthy tissue damage during radiotherapy can be reduced using helium ions [4], which can be further optimized using ultra high dose rate delivery (FLASH). Healthy tissue sparing using ultra high-dose rate deliveries has been predominantly known for electrons and, more recently, proton beams [5–8]. Considering helium ions, two studies have shown that ultra-high dose rate helium beams have significant potential for FLASH sparing effect in vitro [9] and in vivo in a zebrafish model [10]. The best of our knowledge, this is the first study describing FLASH effect with raster scanning helium ions in a preclinical mouse model. In the current study, we investigated the effects of FLASH irradiation, focusing on the brain endothelium and microglia/macrophages. These two components are encompassed in neurovascular unit (NVU) [11] and play a critical role in maintaining brain homeostasis [12]. Endothelial cells within the blood brain barrier (BBB) and NVU express CD31, which can serve as an indicator of endothelial BBB integrity [13]. Reduced CD31 signal may imply BBB/endothelium compromise [14], which is potentially linked to irradiation injury with microglia activation [15]. The activation of microglia, which are the primary immune cells in the brain [16], plays a crucial role in the brain’s response to radiation [15]. Microglia can reside in resting state or activated state. Whereas resting state is characterized by ramified cellular morphology, activation of microglia is followed by a change to ameboid morphology [17, 18]. It is well-established that irradiation induced microglial activation can lead to neural tissue toxicity [19], trigger a potentially damaging neuroinflammatory response [20], and contribute to cognitive impairment [21]. Among various markers of microglial activation, a general phagocytic marker of activated microglia-CD68 [22] has been found to be upregulated in response to helium ion irradiation induced stress [23]. Another marker of microglial response to stress stimuli is increased expression of inducible nitric oxide synthase (iNOS) [24, 25]. Nitric oxide (NO) is a regulatory molecule that play a role in regulating brain homeostasis under physiological conditions [26]. Introducing a novel radiotherapy approach necessitates demonstrating its efficacy in maintaining tumor control equivalent to standard treatments. Previous studies have shown that FLASH electron and proton [27–32], as well as carbon ion beams [33] effectively eradicate tumors, subsequently broadening the therapeutic window. In this study, we present the first outcomes on tumor control following high-dose rate helium ion treatment. The primary objective of this study was to investigate the potential of raster-scanned helium ions to achieve the FLASH effect by examining the response in healthy brain tissue and assessing their efficacy in tumor control.

## Materials and Methods

### Planning and irradiation

The irradiation setups were located near the beam exit and included a 2D ridge filter. The 2D ridge filters were used to create a 2 cm spread out Bragg peak (SOBP) for the brain irradiation and a 1 cm SOBP for tumor irradiation, respectively. PMMA absorbers (112 mm for the brain and 114 mm for the tumor) have been included in the setup as previously described [9]. The initial beam energy was 145.74 MeV/u.

For the brain irradiation, treatment plan was consisting of 16 spots, with a 4 mm spot spacing at isocenter, resulting in a 12×12 mm^2^ field. The plan was optimized to deliver 10 Gy in the center of each brain for a maximum dept of ~6 mm, for both SDR and FLASH irradiations. For SDR brain irradiations, the delivery of the mono-energetic layer was divided into 10 spills (instant spill dose rate: 2.6 Gy/s; spill time: 0.38 s, delivery time: 48 s) to achieve an average dose rate closer to clinical scenarios (~0.2 Gy/s). FLASH dose rate was 141 Gy/s in one spill (spill time 0.07 s). A collimator (material: PMMA; thickness 1cm, aperture: 6 × 6 mm^2^) was placed on top of the mouse heads. For the tumor irradiation, treatment plan was consisting of 25 spots, with a 2 mm spot spacing at isocenter, resulting in an 8×8 mm^2^ field. The plan was optimized to deliver 12.5 Gy over the entire tumor for both SDR and FLASH irradiations. For SDR tumor irradiations, the delivery of the mono-energetic layer was divided into 8 spills to achieve an average dose of ~0.2 Gy/s (instant spill dose-rate 0.4 Gy/s; spill time 4 s, delivery time 60 s). FLASH irradiation was delivered at 250 Gy/s to the tumor site in one spill (spill time: 0.05 s)

For in silico verification and physical distribution quantification, a detailed geometry of the in vivo experimental set-up was incorporated into the in-house FLUKA Monte Carlo (MC) simulation system, including specifications of the HIT beam-line. Simulations were performed for dose and dose-averaged linear energy transfer (LET_D_) prediction. For both delivery cases, measurements with ionization chamber (PinPoint chamber 31015, PTW) following TRS398 were performed to verify the dose at the desired position, with and without collimator. Additional verifications were performed with EBT gafchromic films.

### Animal model

Six-week-old female C57BL/6 mice (Janvier Labs; RRID: IMSR_RJ:C57BL-6JRJ) were used in this study. Mice were anesthetized with isoflurane for irradiation and fixed in a custom-made holder during irradiation. A collimator was placed over the head of each mouse, sparing an area of 6 × 6 mm^2^ above the right hemisphere as the irradiation field. Animal work was carried out in accordance with the rules approved by the local and governmental animal care committee established by the German government (Regierungspraesidium, Karlsruhe).

For establishing a subcutaneous carcinoma tumor model, adenocarcinoma cells A549 (purchased from ATCC; RRID: CVCL_0023) were used and were confirmed to be Mycoplasma negative by PCR. Cells were cultured in Dulbeccós Modified Eagle Medium (DMEM, ATCC) supplemented with 10% heat-inactivated Fetal Bovine Serum (FBS, Millipore), 2 mM glutamine and 1% Penicillin/Streptomycin (Gibco) at 37°C at 5% CO_2_ atmosphere. 5×10^6^ A549 cells were resuspended in 100 µl PBS and injected subcutaneously in a hinder limb of 4-5 weeks old female NMRI mice (Rj: NMRI-Foxn1 nu/nu; Janvier Labs; RRID: IMSR_RJ: NMRI-NUDE). Mice were randomized for irradiation when the tumor volume reached 80 ± 20 mm^3^. For irradiation, mice were positioned in the holders and subcutaneous tumors were irradiated using 8×8 mm^2^ field size, assuring complete tumor target coverage. Tumor sizes were measure 2-3 times per week using a caliper and volumes were calculated with the following formula: Tumor volume (mm^3^) = length × width × width/2. To compare the percentage survival between different treatment groups, Kaplan-Meier survival curves were generated.

### Tissue preparation and immunofluorescence

Tissue was prepared and stained as described [34]. Briefly, mice were anesthetized with 120 mg/kg ketamine (Ketaset, Zoetis) and 20 mg/kg xylazin (Rompun, Bayer) and transcardially perfused with ice-cold PBS (Gibco) followed by ice-cold 4% PFA (Carl Roth) in PBS. The mice were decapitated and brains were fixed in 4% paraformaldehyde and cryoprotected with 30% sucrose (Roth) in PBS until they sank. Brains were placed in Tissue-Tek Cryomolds (Sakura Finetek) and embedded in Tissue-Tek O.C.T. Compound (Sakura Finetek) before slow freezing in isopentane (2-Methylbutane, Honeywell; pre-cooled in liquid nitrogen) and subsequent storage at −80°C. Brains were sagittally sectioned (thickness 8–10 µm) on a cryotome (CryoStar NX70; Thermo Scientific). Sections were thawed at room temperature, washed, and blocked using blocking buffer containing 2.5% goat serum, 1% bovine serum albumin (BSA), and 0.1% Triton X in phosphate-buffered saline (phosphate-buffered saline (PBS)) and ReadyProbesTM Mouse-on-Mouse IgG Blocking Solution (Invitrogen, Carlsbad, CA, USA) according to the manufacturer’s instructions. CD31 (1:50; Abcam Cat# ab28364, RRID: AB_726362) antibody was used for endothelial cell and blood vessel assessment. Iba1 (1:500; FUJIFILM Wako Pure Chemical Corporation Cat# 019-19741, RRID: AB_839504) and CD68 antibody (1:100, Bio-Rad Cat# MCA1957GA, RRID: AB_324217) were used for microglia/macrophage identification. iNOS was used as a marker of microglia activation (1:100; Abcam Cat# ab3523, RRID: AB_303872). Slides were washed and signals were detected using Alexa Fluor antibodies (for γH2AX: Thermo Fisher Scientific Cat# A-21236, RRID: AB_2535805; for Iba1 and iNOS: Thermo Fisher Scientific Cat# A-21206, RRID: AB_2535792; for CD68: Thermo Fisher Scientific Cat# A-21247, RRID: AB_141778; for CD31: Thermo Fisher Scientific Cat# A-21428, RRID: AB_2535849) as secondary antibodies (1:400). Specificity of staining was confirmed using negative controls where only secondary antibody was applied on the samples. DAPI (1:1000, Sigma Aldrich) dye was used to stain nuclei.

### Microscopy and image analysis

Zeiss Axio Scan.Z1 microscope equipped with a Colibri 5/7 with 385 nm, 567 nm, and 630 nm LEDs, 20x 0.8 Plan-Apo objective and Orca Flash 4.0 V2 sCMOS camera was used for imaging. The images were then analyzed using the high throughput software ScanR Analysis (v3.2) and ImageJ (RRID:SCR_003070). For γH2AX, a nuclei detection based on DAPI intensity thresholding was performed followed by a sub-object routine which detected nuclear γH2AX signal above a defined fixed threshold based on the control (non-irradiated) samples. The resulting nuclei which were positive for DAPI and γH2AX were scored. For γH2AX nuclear signal total intensity analysis, total intensities within control samples were averaged and normalized to 1, and total intensity signals from the treated cells were normalized to the averaged control signal. The change in γH2AX total intensity signal was then expressed as a fold change in comparison to the control. For CD31 detection a thresholding and edge detection method was utilized to detect the CD31 positive blood vessels. CD31^+^ microvascular density was calculated as the proportion of CD31^+^ signal area over the total measurement ROI area and expressed as a percentage.

To measure and quantify the percentage of Iba1, CD68 and γH2AX positive cells, in-house developed macros for ImageJ were used. In short, images of DAPI were background subtracted using Rolling Ball algorithm, blurred with Gaussian blur, segmented with Find Maxima tool and thresholded. The output of Find Maxima was Single Points (centers of DAPI stained nuclei). The images of stained area were processed with Rolling Ball Background Subtraction, Gaussian Blur and the selection was created above threshold. All single points (centers of DAPI) within the selected stained area were counted and compared to the total cell count. Number of positive stained cells was normalized to the total number of nuclei. For all images, analyses were performed in irradiated cortical brain area in order to keep structural consistency of the investigated brain region among the samples. Each tissue was analyzed using 12-15 ROIs (each ROI equaled 1000×1000 pixels). Five brains per group were analyzed, except for γH2AX analysis, where three brains per group and per time point were analyzed.

### Statistical analysis

Data presentation and statistics were done in GraphPad Prism software (RRID:SCR_002798). One-way analysis of variance (ANOVA) with Tukey’s multiple comparisons test was used to compare the treatment outcomes. Log-rank test was used to compare Kaplan-Meier survival curves.

### Data Availability

The data generated in this study are available upon request from the corresponding author.

## Results

### Irradiation parameters and settings

Helium ion beams at SDR and FLASH dose rates were delivered as described in Material and Methods. For healthy brain irradiation, experiment schematic is displayed in Fig.1a, whereas profiles of dose and LET_D_ as function of depth within the mice brain are shown in Fig.1 b and c. LET_D_ range in high dose region was from 20 keV µm^-1^ to 40 keV µm^-1^. Subcutaneous A549 tumors (Fig. 1d) were subjected to irradiation using 12.5 Gy helium ions for both SDR and FLASH dose rates in order to achieve tumor control. Profiles of dose and LET_D_ as function of depth within the tumor are shown in Fig.1e and Fig.1f. LET_D_ range in the tumor region was from 14 keV µm^-1^to 37 keV µm^-1^.

**Figure 1.**
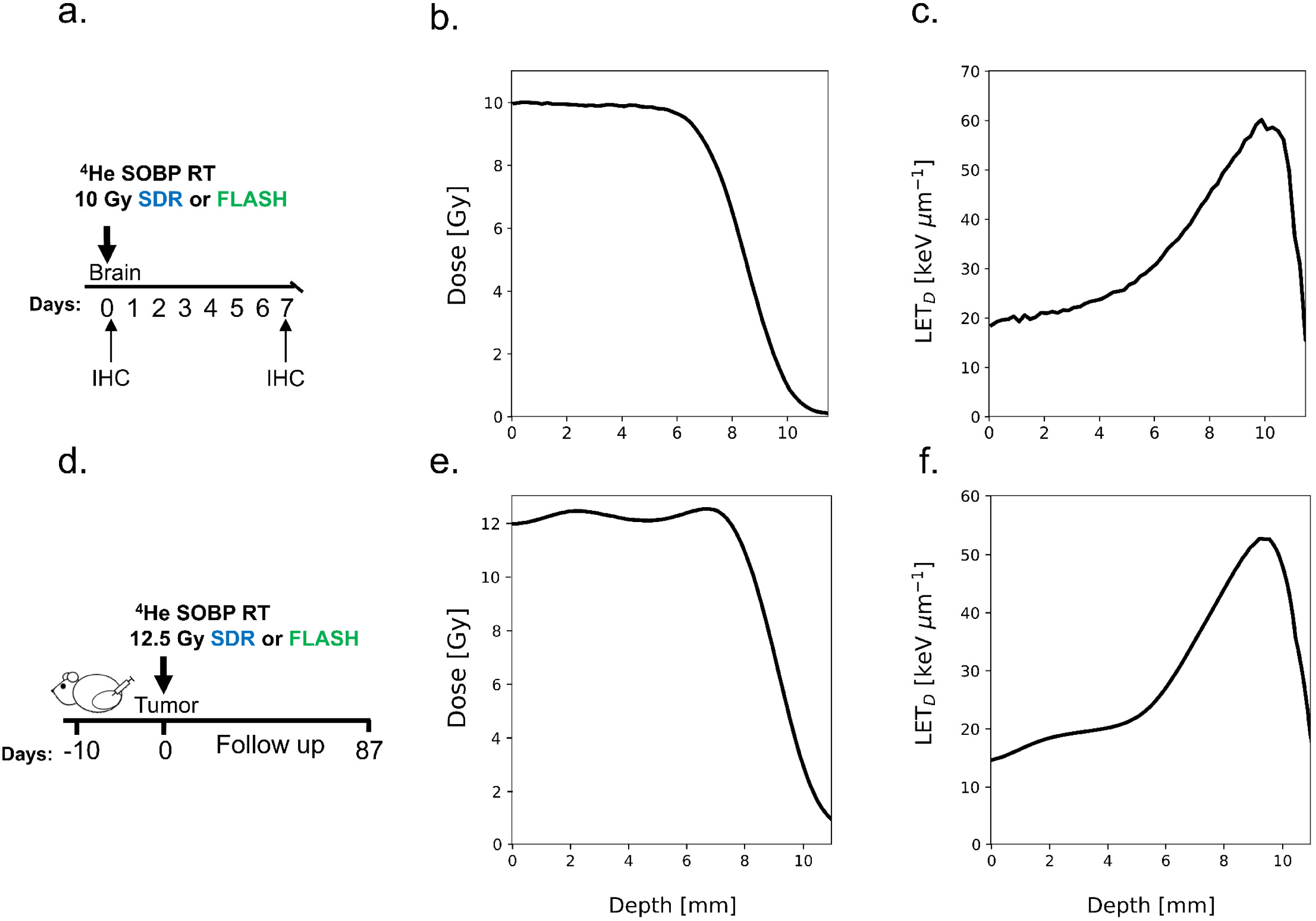
Irradiation parameters. **a.** Experimental scheme for healthy brain irradiation. **b.** Monte Carlo simulated depth dose distributions of the 10 Gy SOBP. **c.** Monte Carlo simulated depth dose-avereged LET (LET_D_) distributions of the 10 Gy SOBP. **d.** Experimental scheme for s.c. tumor irradiation. **e.** Monte Carlo simulated depth dose distributions of the 12.5 Gy SOBP. Monte Carlo simulated depth dose-avereged LET distributions of the 12.5 Gy SOBP.

### FLASH RT spares brain tissue from DNA damage

To investigate DNA damage in brain tissue, after SDR vs. FLASH helium radiotherapy, we used γH2AX as a double-strand DNA break (DSB) marker. The image analysis indicated significantly reduced intensity of γH2AX nuclear signal after FLASH irradiation, at both early (1h-post RT; p= 0.0083; Fig. 2a) and late (7 days post-RT, p= 0.0001; Fig. 2b) time points. Moreover, the fraction of γH2AX positive nuclei was significantly lower in FLASH-irradiated samples than in SDR-irradiated samples (p= 0.0001), which could indicate for differential double-strand breaks damage formation and/or repair in FLASH vs. SDR RT (Figure 2).

**Figure 2.**
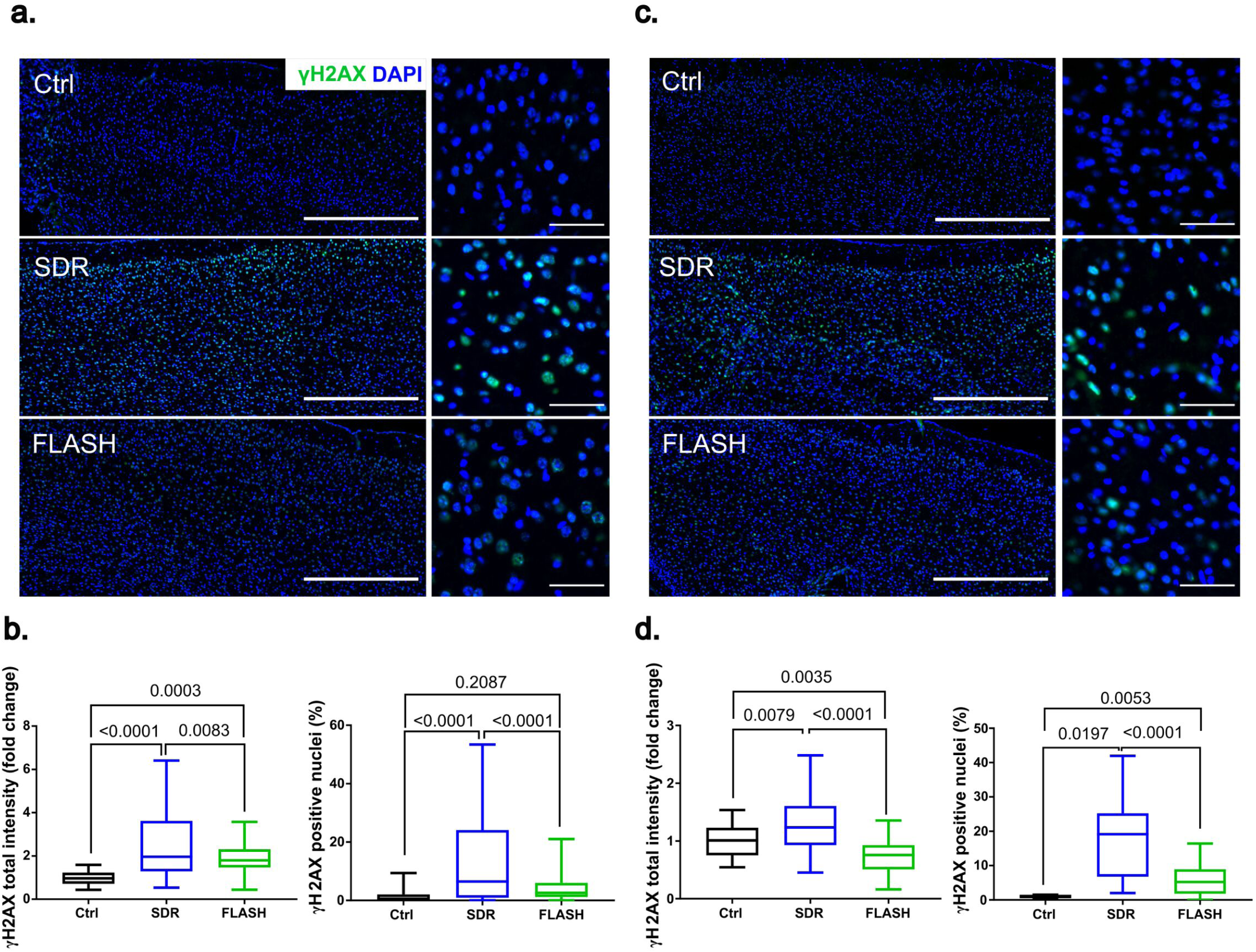
FLASH RT results in lower pS139 H2AX foci (γH2AX) signal in healthy C57BL/6 mouse brain sections. **a.** Representative images and **b.** quantification of γH2AX total intensity (shown as a fold change normalized to the non-irradiated control) and percentage of γH2AX positive nuclei at 1h post-irradiation (N_Ctrl_=3, N_SDR_= 3, N_FLASH_= 3 brains). **c.** Representative images and **d.** quantification γH2AX total intensity and percentage of γH2AX positive nuclei at 1 7 days N_Ctrl_=3, N_SDR_= 3, N_FLASH_= 3, brains) post-RT. Arrows point to the γH2AX positive nuclei. Non-irradiated samples were used as controls. Data are presented as box and whiskers plots (min, max; horizontal line indicates median). Groups were compared using ANOVA Tukey’s post-hoc test. (overview scale bar: 500 µm; zoom scale bar: 50 µm; magnification 20x).

### FLASH beams preserve brain microvascular architecture

Structural changes of the CD31^+^ endothelium were examined to assess the integrity of brain tissue after irradiation. As shown in Fig. 3a, the CD31 signal of the brain microvascular endothelium is more evident for control samples, than in irradiated samples (7 days post-RT). Quantification of CD31^+^ structures indicated significant decrease in CD31 microvascular density (MVD) after radiotherapy (p= 0.0001, Fig 3b). However, the radiation-induced change in MVD was dose rate-dependent. Brains irradiated with FLASH beams exhibit higher CD31^+^ MVD in comparison to SDR irradiated tissues (p= 0.018, Fig. 3b).

**Figure 3.**
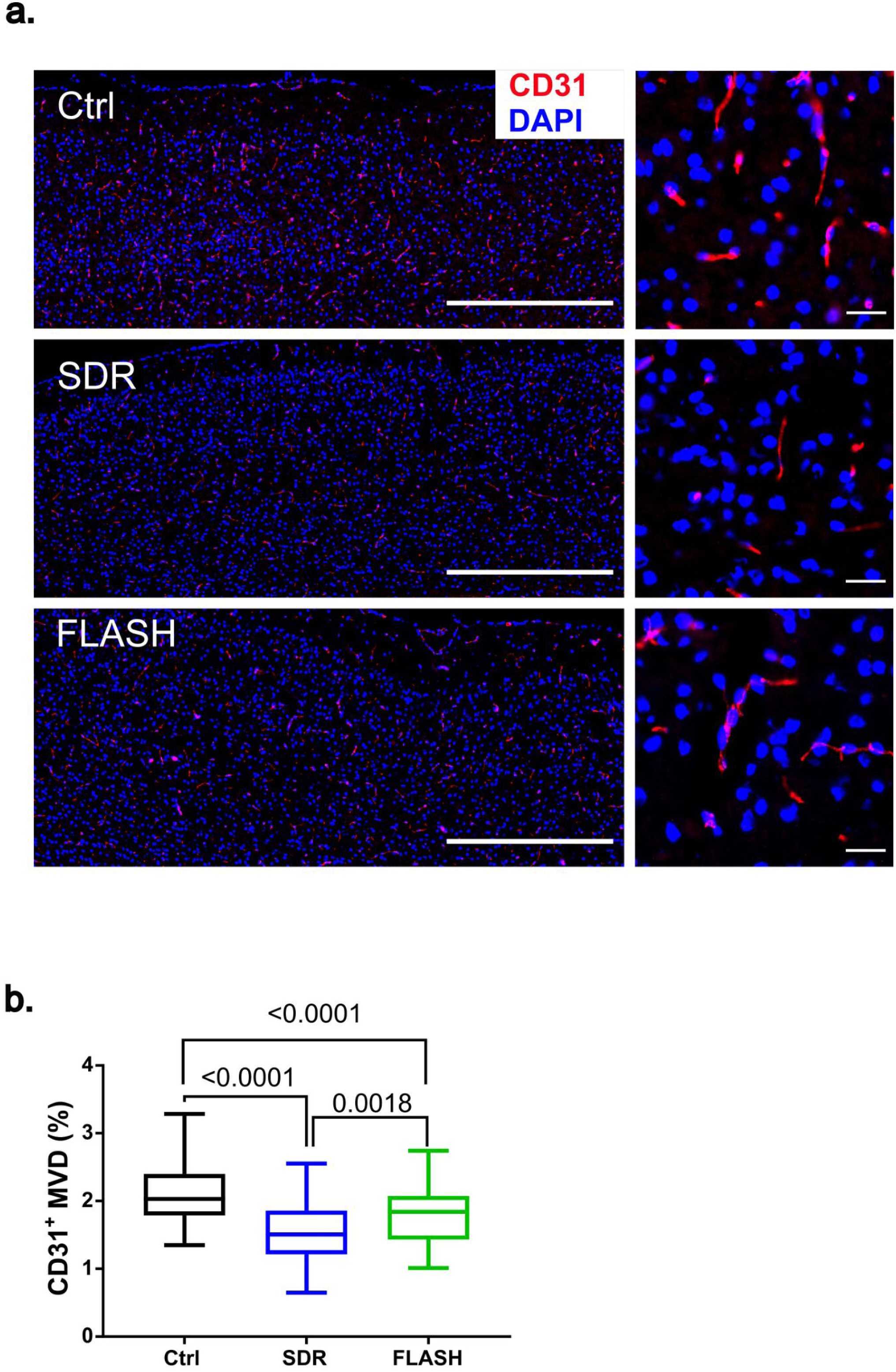
FLASH beams spare brain microvascular architecture. **a.** Representative images of CD31 stained brain sections 7 days post-RT (overview scale bar: 500 µm; zoom scale bar: 25 µm; magnification 20x). **b.** CD31^+^ microvascular density (MVD) was shown for each condition (N_Ctrl_= 5, N_SDR_= 5, N_FLASH_= 5 brains). Data are presented as box and whiskers plots (min, max; horizontal line represents median). Groups were compared using ANOVA Tukey’s post-hoc test.

### FLASH RT exhibits reduced microglia/macrophages activation

Both SDR and FLASH RT decreased the ramified (surveilling [35]) Iba1^+^ microglia/macrophage population (Fig. 4). Considering the morphology of Iba1^+^ microglia/macrophages after radiotherapy, we could observe a more prominent rounded cell shape in treated groups (Fig. 4a), which is commonly associated to the activated phenotype [36]. Images of the tissue sections in the FLASH group indicated mixed presence of both ramified and round microglia, but this observation could not be statistically confirmed.

**Figure 4.**
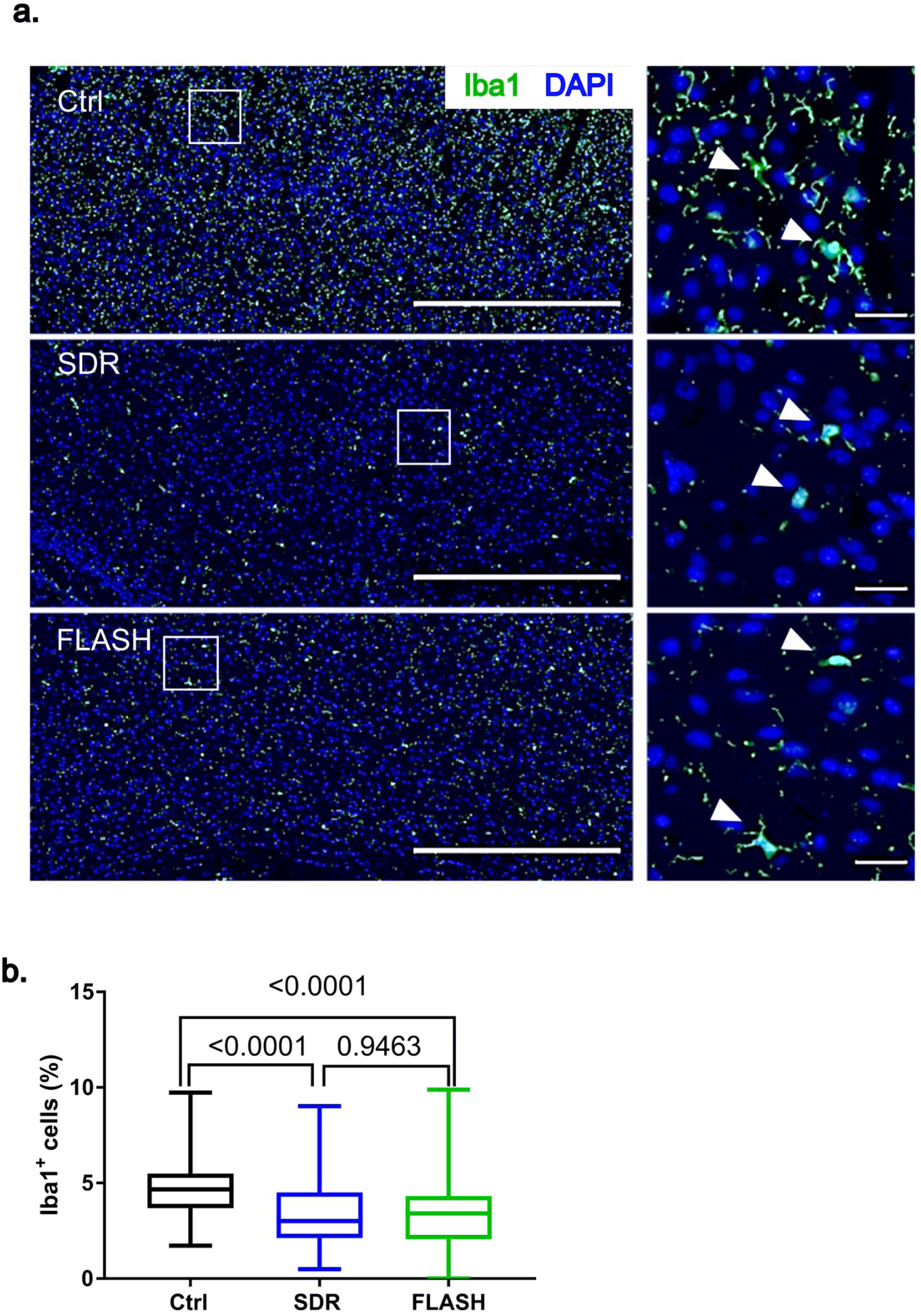
Radiotherapy decreases levels of ramified microglia. **a**. Representative images of Iba1^+^ microglia 7 days post-irradiation of the mouse brain cortex (overview scale bar: 500 µm; zoom scale bar: 25 µm; magnification 20x). Arrow heads indicate ramified microglia in Ctrl group, and more rounded microglial morphology in SDR and FLASH groups. **b.** Quantification of Iba1^+^ cells. Data are presented as box and whiskers plots (min, max; horizontal line represents median; N_Ctrl_= 5, N_SDR_= 5, N_FLASH_= 5 brains). Groups were compared using ANOVA Tukey’s post-hoc test.

SDR RT induced a significant increase in the cellular fraction of CD68^+^ microglia/macrophages (p= 0.0001). FLASH RT and control samples showed a similar percentage of the CD68^+^ population (Fig. 5b). CD68^+^ cells were characterized by an ameboid shape (Fig. 5a), which is representative of activated, phagocytic microglia/macrophages [24]. Another hallmark of microglia/macrophages activation is an increase in iNOS expressing cell population, which could be observed after SDR RT (p= 0.0268; Fig. 5c). iNOS expression was evident in most of CD68^+^ cells (Fig. 5a). SDR treatment led to an increase of this double-stained population when compared to the control (p= 0.0002) or FLASH irradiated tissues (p= 0.0038). Control and FLASH samples were characterized by similarly low levels of CD68^+^iNOS^+^ cells (Fig. 5d).

**Figure 5.**
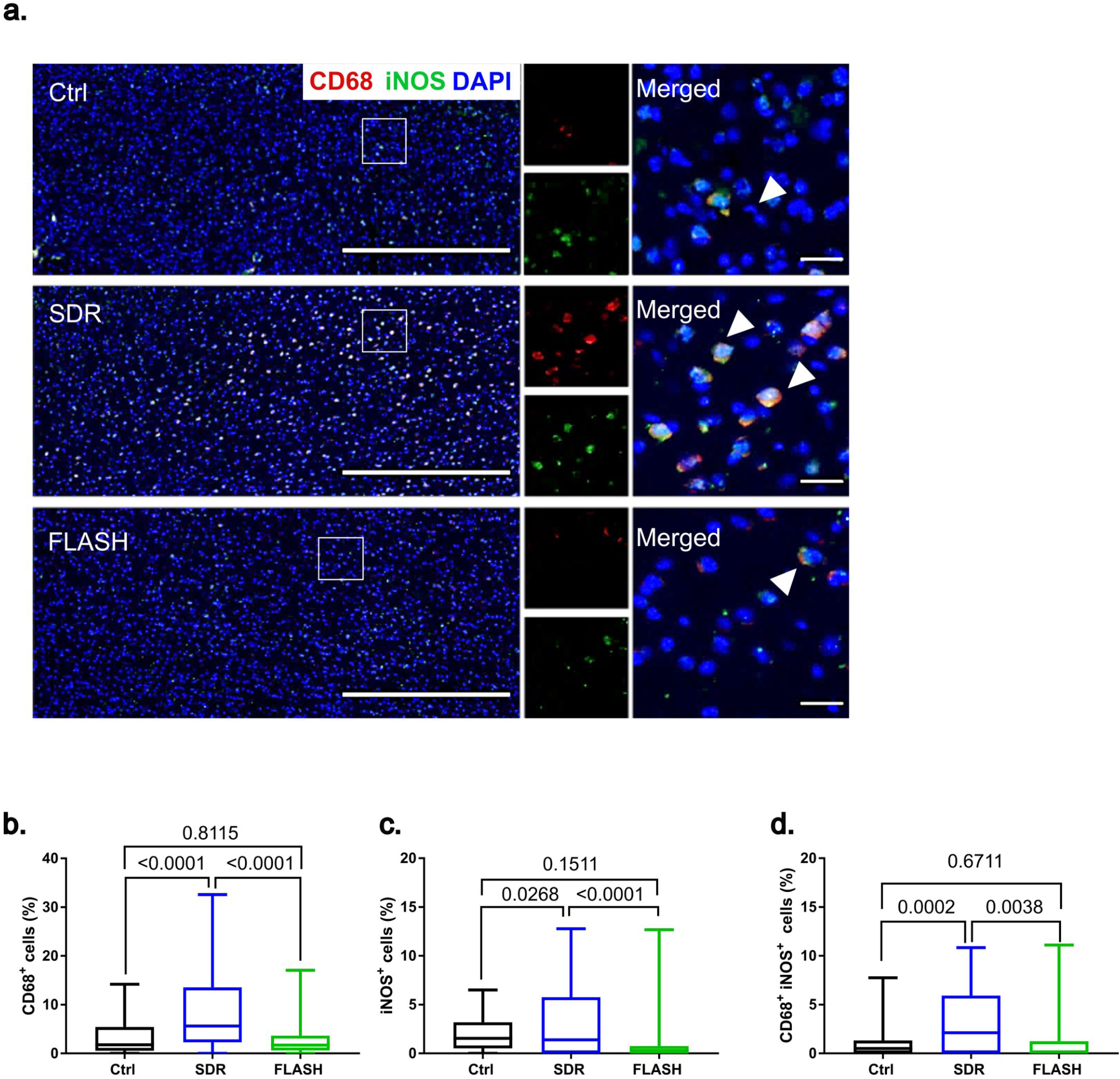
FLASH radiotherapy results in reduced phagocytic microglia/macrophages activation. **a**. Representative images of CD68^+^ and iNOS^+^microglia/macrophages of the mouse brain cortex 7 days post-irradiation (overview scale bar: 500 µm; zoom scale bar: 25 µm; magnification 20x). Arrow heads indicate CD68 and iNOS double positive cells. Quantification of **b.** CD68^+^, **c.** iNOS^+^ and **d.** CD68^+^iNOS^+^ co-stained cells. Data are presented as box and whiskers plots (min, max; horizontal line represents median; N_Ctrl_= 5, N_SDR_= 5, N_FLASH_= 5 brains). Groups were compared using ANOVA Tukey’s post-hoc test.

### FLASH RT preserves tumor control

In this model, both SDR and FLASH radiotherapy significantly reduced tumor volume compared to control mice 14 days post-treatment (p< 0.0001; Fig. 6d). Both modalities also significantly prolonged survival relative to the control group, with median survival times of 24 days for the SDR group (p= 0.012) and 43 days for the FLASH group (p= 0.003), compared to 14 days for the control group (Fig. 6c). While animals treated with FLASH survived longer than those treated with SDR, this difference was not statistically significant (p= 0.076).

**Figure 6.**
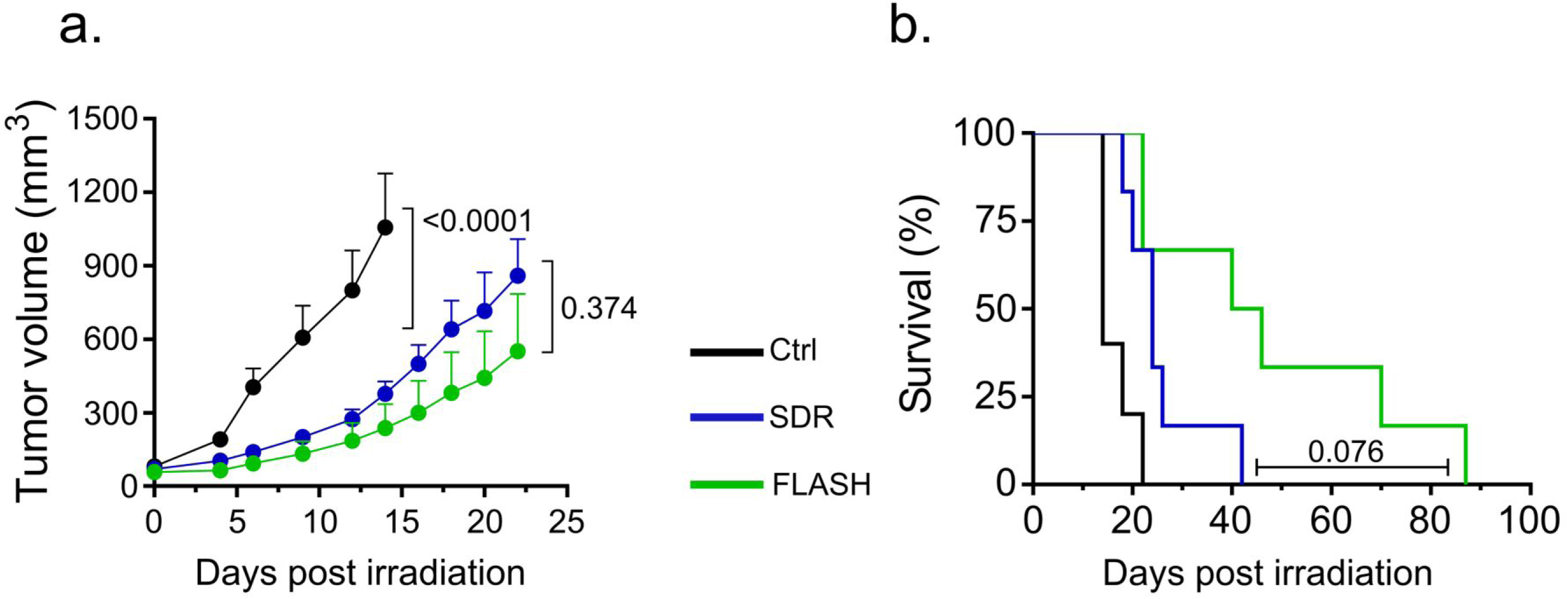
FLASH radiotherapy preserves tumor control **a.** Tumor volumes are presented as the mean ± SEM. The p values were calculated by two-way ANOVA with Tukey’s post-test (n_Ctrl_= 5; n_SDR_= 6; n_FLASH_= 6). **b.** Kaplan-Meier plot of survival by treatment group (Log-Rank test curve comparisons: Ctrl vs. SDR p= 0.012; Ctrl vs. FLASH p= 0.003; SDR vs. FLASH p= 0.076; N_Ctrl_= 5; N_SDR_= 6; N_FLASH_= 6).

## Discussion

Helium ion radiotherapy may offer advantages over conventional photon and proton therapies by minimizing damage to healthy tissues and providing comparable target coverage [37]. Combining helium ion irradiation with FLASH dose rate delivery may emerge as the therapy of choice for cases requiring higher LET and exceptional normal tissue sparing.

In this study, the FLASH effect of helium ion radiotherapy was examined in preclinical settings in vivo. For this purpose, healthy mouse cortex was irradiated with 10 Gy helium ions using either ultra-high dose rate (FLASH) or standard dose rate (SDR). We compared the impact of helium FLASH radiation to SDR radiation on brain toxicity focusing on two aspects: the formation of DNA double-strand breaks (DSB) and the structural integrity of the neurovascular unit in the cerebral cortex (the microvascular endothelium and the activation of microglia/macrophages). In comparison to SDR, FLASH radiotherapy led to the reduced accumulation of the DSB DNA-damage which was visualized using DSB surrogate marker γH2AX [38]. γH2AX signal intensity, as well as γH2AX positive cell nuclei, were significantly lower in the FLASH group. The reduction of γH2AX signal was apparent both at 1 hour and 7 days post-RT. Reduced DNA damage after FLASH therapy is consistent with previous in vivo observations after proton FLASH irradiation [6, 27], in vitro using ultra-high dose rate helium beams in low oxygen condition [9] and ex-vivo using ultra-high dose rate electrons [39].

Irradiation-induced brain injury has been linked to microvascular endothelial damage, glial cell destruction, and inflammation, leading to a disruption of the neurovascular unit [40]. These adverse effects on brain tissue have been previously reported for helium irradiation [23, 41]. An increase in CD68^+^ microglia and an upregulation of iNOS after SDR therapy in this work, aligns with existing literature that suggests irradiation-mediated microglial activation [42]. The co-expression of CD68 and iNOS is frequently associated with pro-inflammatory and phagocytic microglia/macrophages [25]. Here we demonstrate that at least a part of these side effects can be mitigated when helium ions are delivered at FLASH dose rate, similarly to what was previously shown for FLASH electrons [8]. SDR treatment led to an increase in activated microglia/macrophages (CD68^+^), while significantly reducing CD31^+^ endothelial structures. In contrast, FLASH treatment had minimal impact on the CD68^+^ population but moderately reduced CD31^+^ structures compared to control. This is in line with a previous study showcasing neuroprotective effects of proton FLASH beams [34] and complementary to the literature indicating electrons FLASH sparing of CNS vasculature [43] and neurocognitive function [8, 44]. Interestingly, both SDR and FLASH RT decreased Iba1^+^ ramified microglia (also called surveilling microglia) morphology, often associated with their physiological state [28, 45]. Even though several studies have reported a reduction in Iba1 expression following radiotherapy [46–48], the complex role of microglial activation following irradiation injury remains only partially deciphered [15]. However, transition of microglia to a pro-inflammatory state may lead to neural and endothelial tissue damage [49]. Microglia interacts with endothelial cells to maintain a stable brain environment and regulate BBB integrity [25]. Dysregulation of these interactions can lead to neurological diseases, emphasizing the importance of the microglia-endothelium connection in brain health. Reduced microglial transition to the pro-inflammatory phenotype after FLASH irradiation, together with preservation of the CD31^+^ microvascular endothelium, indicates benefits of FLASH helium ions in preserving brain microenvironment.

In addition to its beneficial effects on healthy tissue, helium ion FLASH radiotherapy achieved tumor control comparable to SDR treatment, further confirming the potential of FLASH helium ions as a promising therapeutic modality. To the best of our knowledge, this is the first report providing evidence of the FLASH effect of helium ions in mouse tissue, demonstrating effective tumor control relative to SDR radiotherapy. Although the increased survival of FLASH treated mice was not statistically significant, these data suggest the potential superiority of FLASH radiotherapy for tumor treatment, similar to previous findings with FLASH protons [50]. This research lays a groundwork for further exploration of the FLASH effect on the dynamic interaction between brain endothelium and microglia, which are crucial for neuroprotection and functional recovery post-radiotherapy. Future studies should investigate these effects at later stages, including neurocognitive testing and BBB permeability, as well as tumor response including larger cohorts, to explore the long-term benefits of FLASH radiotherapy.

## Conclusion

This work validates the potential of FLASH raster scanning helium ions to induce FLASH sparing effect in cerebral tissue, as compared to standard dose rate in a preclinical in vivo model, while preserving tumor control. Furthermore, this investigation paves the way for additional scrutiny of the ultra-high dose rate effect in particle therapy, which can be relevant in the context of advancing therapeutic strategies in radiation oncology.

## Conflicts of Interest

J.Debus reports grants from CRI – The Clinical Research Institue GmbH grants from View Ray Inc., grants from Accuray International Sarl, grants from Accuray Incorposrated, grants from RaySearch Laboratories AB, grants from Vision RT limited, grants from Merck Serono GmbH, grants from Astellas Pharma GmbH, grants from Astra Zeneca GmbH, grants from Siemens Healthcare GmbH, grants from Merck KGaA Accounts Payable, grants from Solution Akademie GmbH, grants from Ergomed PLC Surrey Research Park, grants from Siemens Healthcare GmbH, grants from Quintiles GmbH, grants from Pharmaceutecal Research Associates GmbH, grants from Boehringer Ingelheim Pharma GmbH Co, grants from PTW-Freiburg Dr. Pychlau GmbH. A. Abdollahi reports grants and other support from Merck KGaA, FibroGen Inc., Bayer, Roche, and Merck Serono and other support from BMS and BioMedX outside the submitted work.

## Acknowledgments

This work was supported by intramural funds from National Center for Tumor diseases (NCT-PRO and Biodose programs) and German Cancer Consortium (DKTK), and in part by National Institutes of Health (NIH-1P01CA257904-01A1) and the collaborative research center of the German research foundation (DFG, Unite, SFB-1389, Project number 404521405). I. Dokic was supported by Olympia-Morata Program. The funders had no role in study design, data collection and analysis, decision to publish or preparation of the manuscript. The authors thank Dr. Stephan Brons and the entire HIT accelerator team for establishing and providing the FLASH ion beam and Dr. Uli Weber from GSI for providing the 2DRM. The authors thank Ms. Lena Vogelbacher for excellent technical support.

## Notes

### Competing Interest Statement

JD reports grants from CRI The Clinical Research Institue GmbH grants from View Ray Inc., grants from Accuray International Sarl, grants from Accuray Incorposrated, grants from RaySearch Laboratories AB, grants from Vision RT limited, grants from Merck Serono GmbH, grants from Astellas Pharma GmbH, grants from Astra Zeneca GmbH, grants from Siemens Healthcare GmbH, grants from Merck KGaA Accounts Payable, grants from Solution Akademie GmbH, grants from Ergomed PLC Surrey Research Park, grants from Siemens Healthcare GmbH, grants from Quintiles GmbH, grants from Pharmaceutecal Research Associates GmbH, grants from Boehringer Ingelheim Pharma GmbH Co, grants from PTW Freiburg Dr. Pychlau GmbH. AA report grants and other from Merck and EMD, grants and other from Fibrogen, other from BMS, other from BioMedX, other from Roche, outside the submitted work.

### Summary of Updates

Figure 1 and Figure 2 have been revised. New Figure 6 included, accordingly Results section was updated. Discussion was updated to include new data on tumor response to radiotherapy introduced in the new figure 6. Table has been removed and information is now contained in material and methods section. Material and methods section has been expanded. An additional co-author included. Manuscript title has been changed.

